# A Tensor Decomposition Uncovers the Effect of Ageing on Muscle and Grip-Load Force Couplings During Grasping

**DOI:** 10.1101/2022.02.25.482040

**Authors:** Chang Ye, Seyed Saman Saboksayr, William Shaw, Rachel O. Coats, Sarah L. Astill, Gonzalo Mateos, Ioannis Delis

## Abstract

Do motor patterns of grasp-to-lift movements change as a result of ageing? Previous studies often relied on simple temporal and kinetic variables to unveil differences caused by ageing, yet their neuromuscular origins remain largely unknown. Here we employed a bimanual grasping protocol with younger and older adults and combined measurements of muscle activity with grip and load forces to provide a window into the neuromuscular strategies underlying effective grasping. We introduced a tensor decomposition to identify patterns of muscle activity and grip-load force ratios while also characterising their temporal profiles and relative activation across object weights and participants of different age groups. This approach extracted the motor components underpinning object grasping across participants. We then probed age-induced changes in these components. A classification analysis revealed two motor components that are differentially recruited between the two age groups. Linear regression analyses further showed that advanced age and poorer manual dexterity can be predicted by the coupled activation of forearm and hand muscles which is associated with high levels of grip force. Our findings suggest that ageing may induce stronger muscle couplings in distal aspects of the upper limbs, and a less economic grasping strategy to overcome age-related decline in manual dexterity.

## Introduction

With advanced age comes decline in motor function [42]. An important daily-life skill required for older adults (OA) is the ability to grasp and lift objects [35, 5, 44]. Several daily tasks, such as carrying heavy objects, require coordination of the two hands (bimanual) [51]. However, the majority of research examining grasping in OA has focused on unimanual control [6] despite the ecological validity of bimanual grasping [32] and its growing use for therapy or rehabilitation (e.g., in stroke [28, 33]).

In unimanual settings, OA have been shown to be slower during the pre-loading and loading phases of the lift [5] and exert higher levels of grip force as well as a larger safety margin when lifting objects [5, 7, 17]. These findings indicate that ageing induces differences in grasping strategies [19]. However, our understanding of why OA adapt their grasping strategies is limited. A potential explanation for this gap in the literature is the lack of computational approach that enables characterising the dynamic interaction between arm and hand muscle activations with grip and load force during grasping [29, 20]. As a result, even when differences in force variables are observed as a result of ageing, their neuromuscular origins remain largely unknown [40].

Thus, in this study we investigate how muscle couplings change with advanced age [34] and how these neuromuscular changes associate with the grasping strategies employed by older adults. Specifically, we a) employ a bimanual grasping protocol to evaluate cross-limb coordination during grasping; and b) combine it with measurements of muscle activity (electromyography - EMG) across the two upper limbs to quantify the underlying motor signals that generate these forces and provide a window into the neuromuscular strategies used to perform an effective grasp [41].

Importantly, to assess the effect of ageing on these signals, both younger (<30 years old) and older (>60 years old) adults (YA and OA respectively) performed this motor task. To also consider the effect of object load, the experimental protocol specified 2 pairs of objects of different weights, henceforth denoted as heavy (H) and light (L), to be grasped and lifted by the participants. Thus, participants executed 10 bimanual grasp-to-lift movements for each pair of objects. Overall, the recordings from this experimental protocol can be represented as 5-way arrays, or tensors [45, 30], indexed by space (EMGs and forces), time (time samples from first contact of the object to end of the object lift), objects (H and L), participants (15 YAs and 15 OAs), and trials (10 for each object). For simplicity, we also consider a 4-way tensor representation of the data after averaging over trials to reduce measurement noise; see also Figure 1(A).

**Figure 1.**
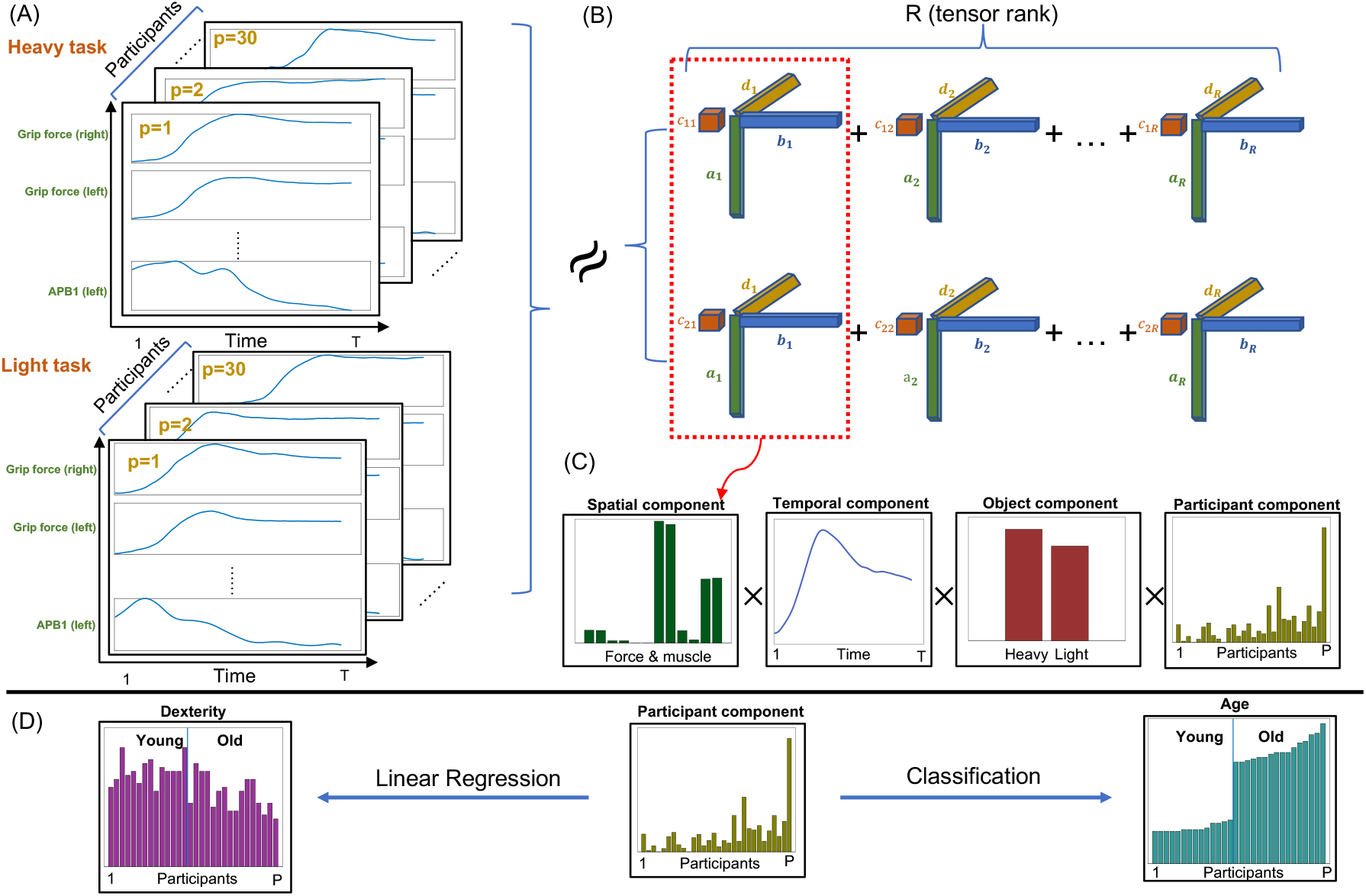
A schematic illustration of the tensor data analysis pipeline implemented in this study. (A) The 4-way tensor data **<u>X</u>** (Spatial×Temporal×Object×Participants); (B) The NCP decomposition **<u>X</u>** ∼ **A** ◦ **B** ◦ **C** ◦ **D**; (C) The first mode of the estimated factor matrices including a spatial (muscle and force) component, a temporal component, an object (heavy or light) component and a participant component; (D) Using the participant component to predict clinical test values (e.g., manual dexterity) or participant characteristics (e.g., age) using regression and classification analyses.

In order to analyze the aforementioned data and unveil the main patterns describing the grasp-to-lift movements across participants, we introduce a high-order (order 5 here, or 4 when signals are averaged across trials) tensor decomposition approach based on non-negative canonical polyadic (NCP, also known as non-negative CANDECOMP/PARAFAC) tensor decomposition; see Figure 1(B). The CANDECOMP/PARAFAC (CP) decomposition is one of the most widespread methods for low-rank tensor factorization, which is unique under very mild conditions [30]. The NCP can also be viewed as an extension of the non-negative matrix factorization (NMF) [31] algorithm typically used to identify muscle synergies from EMG data [49, 8, 1]. The non-negativity constraint of this approach yields a parts-based representation of the underlying motor signals [38]. Crucially, since muscle activations and forces cannot take negative values, the extracted 5-mode components are directly interpretable as muscle activation and force patterns (mode 1) with their corresponding temporal profiles (mode 2) and also specific level of recruitment for each object (mode 3), participant (mode 4) and trial (mode 5); see Figure 1(C) for an example with trial-averaged data and [47, 18, 11] for related approaches with three modes. Said interpretability of the outputs [3, 4] and the lack of any other assumptions, such as orthogonality, enable the extraction of non-orthogonal motor patterns, which are often sparse but partly overlapping, such as the ones typically generated by neural circuits with hard-wired connectivity [53, 37, 9]. Furthermore, the NCP decomposition has only one free parameter, the rank of the tensor *R*, i.e., the number of outer-product components that are sufficient to describe the input data [30]. These properties make the NCP decomposition an attractive and natural method for identifying low-dimensional representations of muscle activations and resulting forces that may differ across time, repetitions, experimental conditions and/or individuals.

Here we ask if motor and kinetic patterns of grasp-to-lift movements change as a result of ageing. We thus use this tensor analysis methodology as a substrate of a machine learning layer to uncover these changes, characterise the functional roles of the underlying signals, and relate them to manual dexterity differences across individuals; see Figure 1(D).

## Results

We firstly fed the EMG and force data recorded during grasp-to-lift movements to a NCP decomposition algorithm to obtain a spatial and temporal characterisation of the main motor and kinetic patterns across participants and objects. Secondly, to investigate the relationship between age and manual dexterity with the activation of each component, we applied linear classification and regression analyses. We first used the estimated participant-mode components (representing component recruitment for each participant) to predict which age group, OA or YA, each participant belongs to. We then employed linear regression and correlation analyses to quantify the contribution of each component to the prediction of age and manual dexterity.

### Unveiling space-time-object-participant patterns of muscle activity via tensor decomposition

The *P* = 30 participants were asked to bimanually grasp and lift heavy (H) or light (L) objects, while the EMG signals of four muscles (Anterior Deltoid, ECR, FCR, APB) as well as kinetic variables, namely the grip force (GF) and load force (LF), were recorded on each upper limb (Left-L, Right-R) as temporal sequences with length of *T*_0_ = 500. Each participant performed *K* = 10 repetitions for each object grasp and lift. An advantage of this experiment was the simultaneous recording of kinetic variables and muscle activations across the two limbs which enabled the study of the relationship between force production and the underlying motor signals over time.

After data pre-processing (see Materials and Methods for details), we obtained the EMG data during the dynamic phase of object lifting (from the point of first contact with the object to the end of the lift) as a 5-way tensor 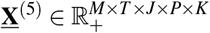, where *T* = 460 is the length of the pre-processed signal sequence; *M* = 12 is the spatial dimension after concatenation of the four forces (Right and Left Grip and Load Forces) and eight EMG signals (from the Right and Left Anterior Deltoid, ECR, FCR and APB muscles) – ordered using the indexing shown in Table 1; *J* = 2 accounts for the heavy (*j* = 1) and light (*j* = 2) objects.

**Table 1.** Indices for the spatial components (mode 1) with descriptions of the measured kinetic variables and EMG-based muscular activity across the two upper limbs.

| Index | Force | Index | Muscle | Index | Muscle |
| --- | --- | --- | --- | --- | --- |
| 1 | grip force (left) | 5 | Anterior Deltoid (left) | 9 | FCR (left) |
| 2 | grip force (right) | 6 | Anterior Deltoid (right) | 10 | FCR (right) |
| 3 | load force (left) | 7 | ECR (left) | 11 | APB (left) |
| 4 | load force (right) | 8 | ECR (right) | 12 | APB (right) |

In order to extract the spatial and temporal components (or factors, both terms will be henceforth used interchangeably), one natural approach is to implement the non-negative CP (NCP) decomposition to the 5-way tensor directly and obtain components in all 5 modes, i.e., spatial, temporal, object, participant, and trial (Figure 2). However, since our main aim here was to characterise age-induced differences across participants, we also considered averaging the measured signals across trials to simplify the data into a 4-way (rather than 5-way) tensor while removing some measurement noise. Before doing this, we assessed the trial-mode output of a 5-way NCP decomposition (see the fifth column in Figure 2 for a *R* = 4 output). We observed minimal variability across trials and, more importantly, no effect of learning/habituation or fatigue in the trial-mode components (which would appear as differential activations between early and late trials). Thus, we reasoned that, for simplicity and reduction of trial-to-trial measurement noise, we could consider the trial-averaged 4-way tensor 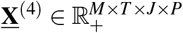.

**Figure 2.**
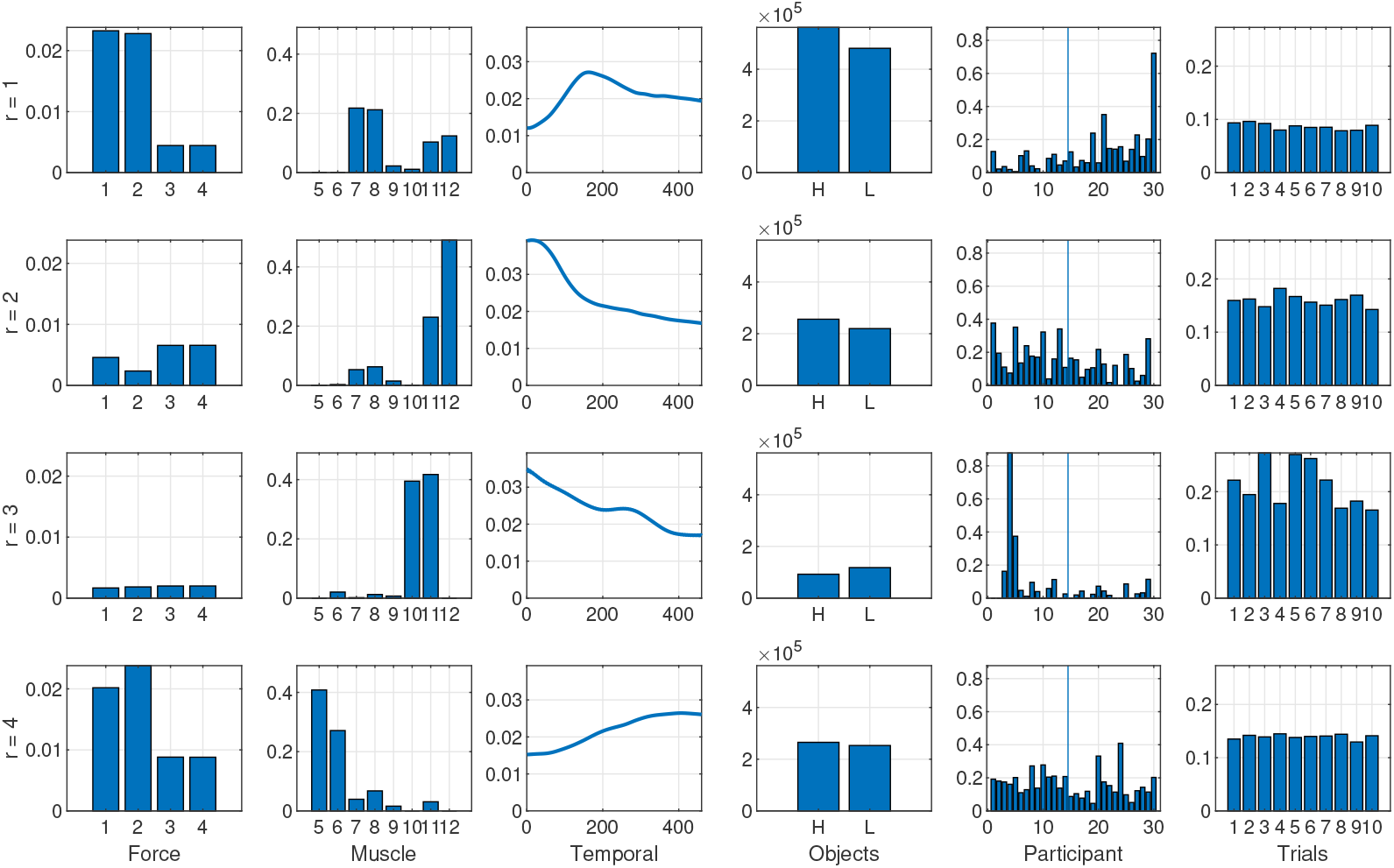
The estimated factors for the 5-way model with *R* = 4 components (in rows). The first and second columns show the decomposition of the spatial mode (*M* = 12) into 4 different components, where the force and muscle modes are separated for improved visualization. Indexing of forces and muscles is done as described in Table 1. The third column shows the four temporal (*T* = 460) components and the fourth column shows the estimated object (*J* = 2) factors. The fifth column shows the participant (*P* = 30) factors, and the vertical line indicates the boundary of the young group (YA, left hand side) and old group (OA, right hand side). The last column shows the four trial (*K* = 10) components.

The next step in our investigation was to select the rank *R* of the NCP decomposition, i.e., how many components suffice to describe the recordings. To this end, we used as criterion a measure of data approximation, namely the Variance Accounted For (VAF, see Materials and Methods for details) and set as criterion the number of components that accounts for *>* 80% of the data variance. We found that *R* = 4 components satisfy this criterion as shown in Figure 4. Figures 2 and 3 show the estimated spatial (force and muscle), temporal, object and participant components of the 5-way and 4-way model. Notice the trial factor in the fifth column of Figure 2. By comparing the *R* = 4 components of the 4-way and 5-way models, it can be seen that the estimated factors are highly consistent (average correlation between spatial factors is *rho* = 0.9967 and between temporal factors *rho* = 0.9981) indicating that the extracted representations are robust regardless of the trial-averaging.

**Figure 3.**
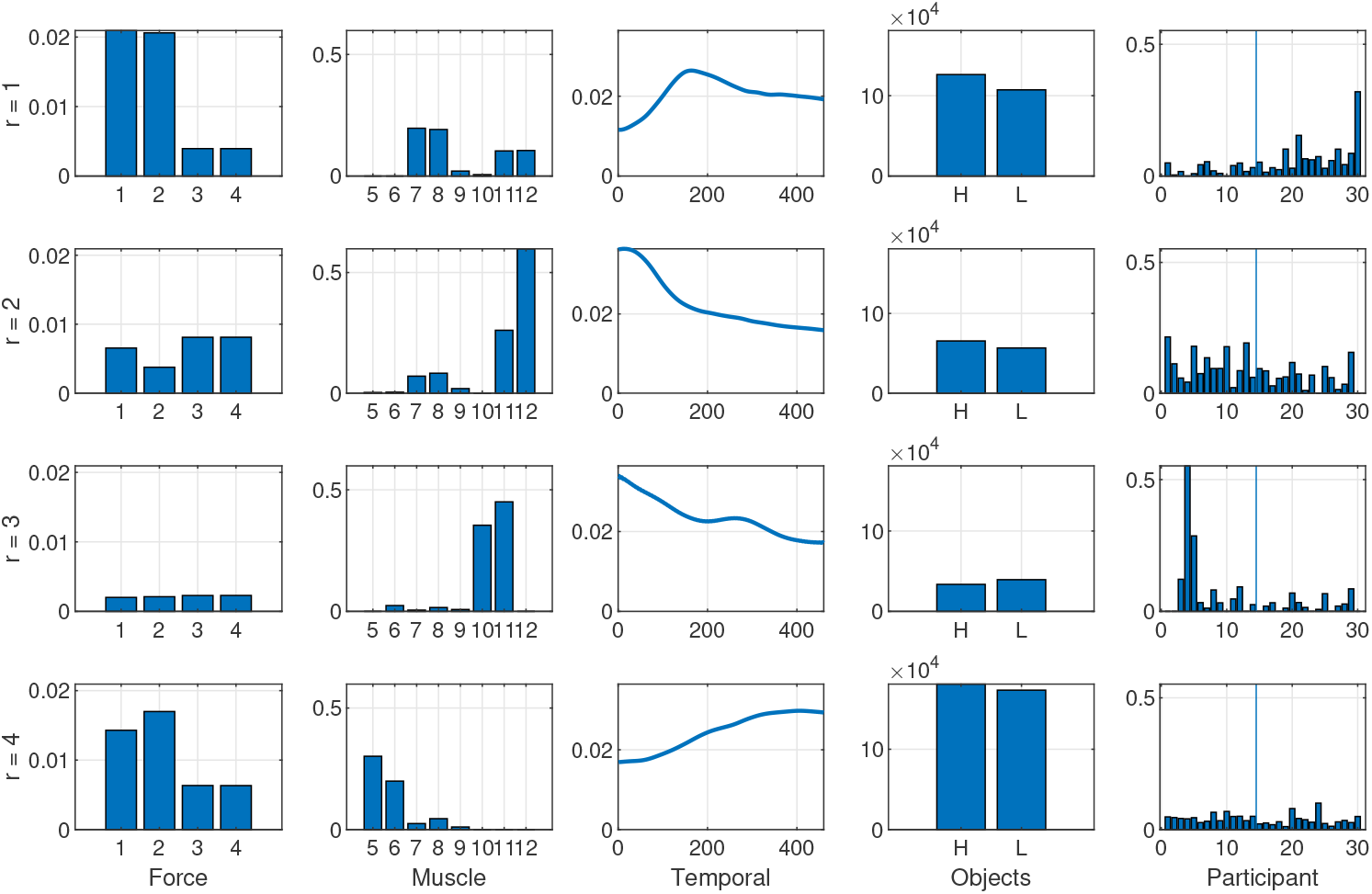
The estimated factors for the 4-way (trial-averaged) model with *R* = 4 components (in rows). The first and second columns show the decomposition of the spatial mode (*M* = 12)into 4 different components, where the force and muscle modes are separated for improved visualization. Indexing of forces and muscles is done as described in Table 1. The third column shows the four temporal (*T* = 460) components and the fourth column shows the estimated object (*J* = 2) factors. The fifth column shows the participant (*P* = 30) factors, and the vertical line indicates the boundary of the young group (YA, left hand side) and old group (OA, right hand side).

**Figure 4.**
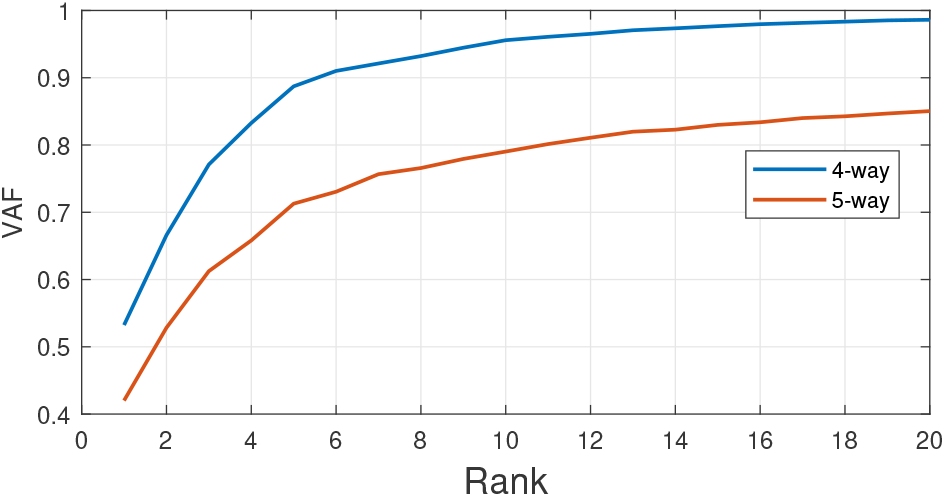
Rank determination for the 4-way and 5-way models. The red and blue curve show the average VAF (10 realizations for each data point) for the 5-way and 4-way models, respectively, obtained for each value of the rank ranging from *R* = 1 up to 20 factors. With *R* = 4 factors, the obtained NCP decomposition of the 4-way data tensor accounts for more than 80% of the variance in the recordings. The data in the 5-way tensor is noisier because trials are not averaged, hence, as expected, the VAF is smaller than its 4-way counterpart for all values of *R*.

The first component (*r* = 1) carries a bimanual grip force representation coupled with bimanual synergies between the extensor carpi radialis longus and the abductor pollicis brevis muscles which synergistically contribute to object grasping and lifting by extending the wrist and abducting the thumb respectively. The participant mode of this factor suggests an increase of the activation of this component with age and the object mode shows recruitment for both object weights with slightly higher activation for the heavy object. The second component (*r* = 2) couples bimanual grip and load forces with high activity of abductor pollicis brevis (primarily the left one) and lower activity of extensor carpi radialis longus and shows a decreasing activation over time and a slight decreasing trend with age. The third component (*r* = 3) captures a co-activation between the left flexor carpi radialis and the right abductor pollicis brevis with a decreasing trend over time, a slightly higher activity for the light object and a sparser recruitment across individuals. This unique pattern probably represents an idiosyncratic motor strategy that is interestingly more representative of light object grasps and lifts. The fourth component (*r* = 4) represents an increasing over time coupling of grip and load forces with bimanual activations of anterior deltoids contributing to arm raising to lift the object.

### Classifying young versus old adults from the identified participant factors

Having characterised the main components of grasp-to-lift movements, we then sought to understand if their recruitment is influenced by the age of the participants. We thus performed a *classification* analysis aiming to predict the age group (OA or YA) of the participant from the participant mode factors.

To identify the most age-discriminating component, we first used each participant factor as a single predictor of age group. We found that the values of the two performance measures (area under ROC curve – AUC and classification accuracy – Acc; see Materials and Methods for details) were (0.82, 0.67), (0.73, 0.67), (0.65, 0.53) and (0.75, 0.70), for components *r* = 1 to *r* = 4 respectively. This indicated that the first component (*r* = 1) had the highest age discrimination power. Then, to identify complementary contributions from other components to age group classification, we considered two or three factors together as predictors for age groups. We found that the highest performance (*AUC* = 0.87, *Acc* = 0.70) was achieved for the combination of the first and second components (Figure 5), suggesting that age group can be best predicted by the joint recruitment of the first two factors (*r* = 1 and *r* = 2).

**Figure 5.**
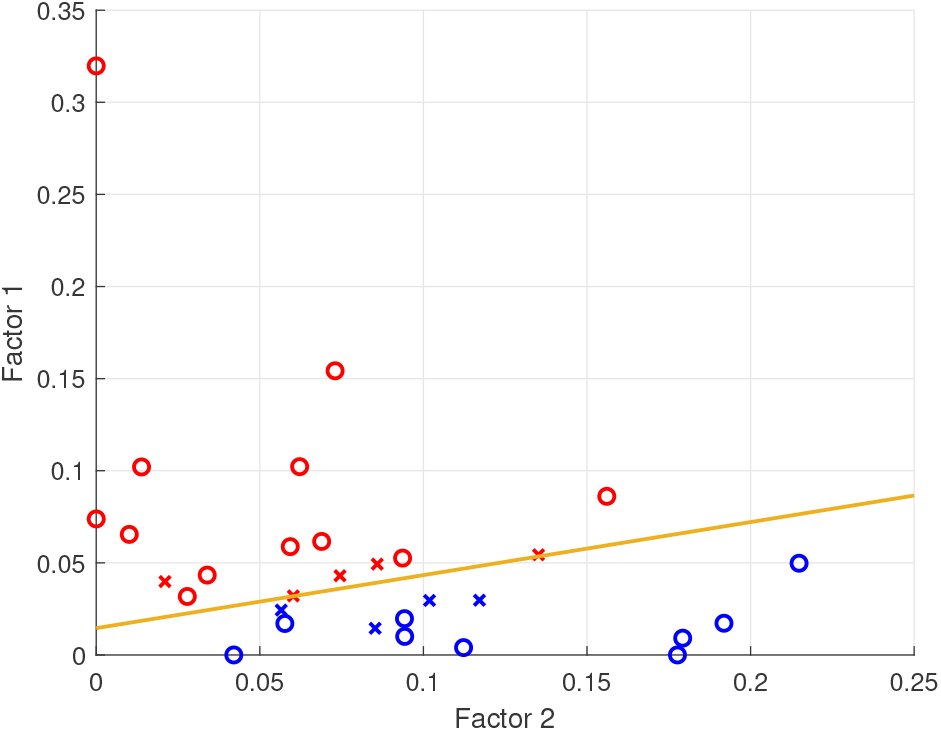
Age group (i.e. YA vs OA) classification using the first two participant-mode factors (*r* = 1 and *r* = 2) of the 4-way model as predictors. Blue and red stand for the predicted labels (blue for YA, red for OA). The circle and cross indicate whether the prediction is correct (circle for correct prediction, cross for wrong prediction). The yellow line indicates the decision boundary of the linear classifier. The AUC and classification accuracy are 0.87 and 0.7, respectively.

In particular, we observed higher (lower) activation of the first component (*r* = 1) for OAs (YAs) and higher (lower) activation of the second component (*r* = 2) for YAs (OAs). This suggests that, although both components were used by all participants regardless of age, their recruitment was dependent on the age group.

### Predicting age and manual dexterity from the identified participant factors

We then asked if the estimated participant factors were predictive of the age and manual dexterity of the participant. We thus considered a *linear regression* model viewing the participant components as the predictors and the age or dexterity vector *PP* as the response. Due to the different normalization procedures for estimated factors of the 4-way and 5-way model, we re-normalized them to unit Frobenius norm (see Materials and Methods for details) prior to linear regression inference. The estimated coefficients of the linear regression model, the p-values of the coefficients and the predictions for different responses (*PP* and age) are depicted in Figure 6 (left and right columns for *PP* and age, respectively). We found that the first component (as well as the intercept term but not the second component) was significantly predictive of both *PP* (*p <* 0.01) and age (*p <* 0.05).

**Figure 6.**
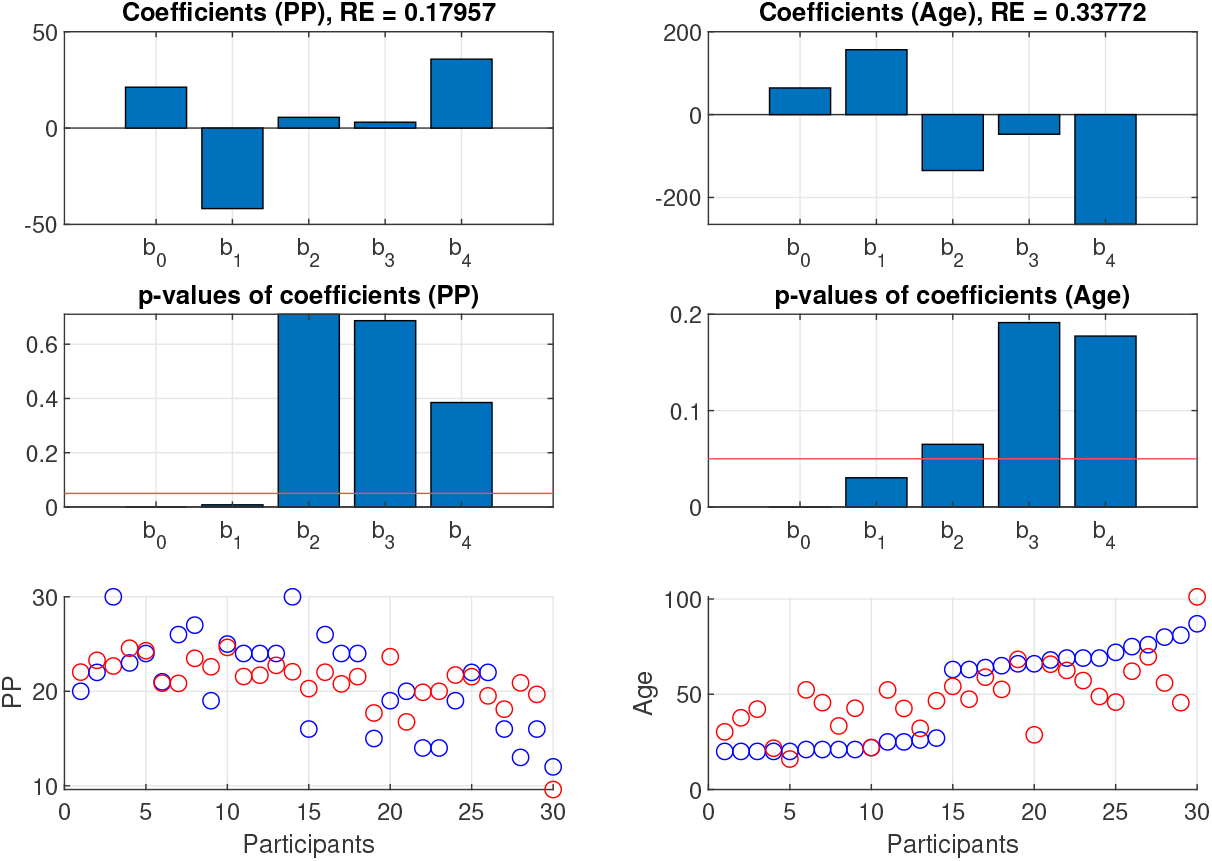
Results of linear regression using the participant modes of the 4-way model as predictors and PP or age as response. The first row shows the regression coefficients, where *b*_0_ is the intercept and *b*_1_, …, *b*_4_ are the regression coefficients for the corresponding participant factors. The second row shows the p-value of the regression coefficients, with the red line indicating significant level 0.05. The third row shows the true and predicted PP (left) or age (right) for different participants. The blue and red points stand for the true and predicted values of PP and age, respectively.

To further probe the (direction of) the relationship between the identified participant factors and age/manual dexterity, we first computed the correlation between the first component (*r* = 1) and the dexterity vector *PP*. We found a significantly negative correlation (*rho* = −0.57, *p* = 0.001) indicating higher activation of the first factor for participants with low dexterity (Figure 7A).

**Figure 7.**
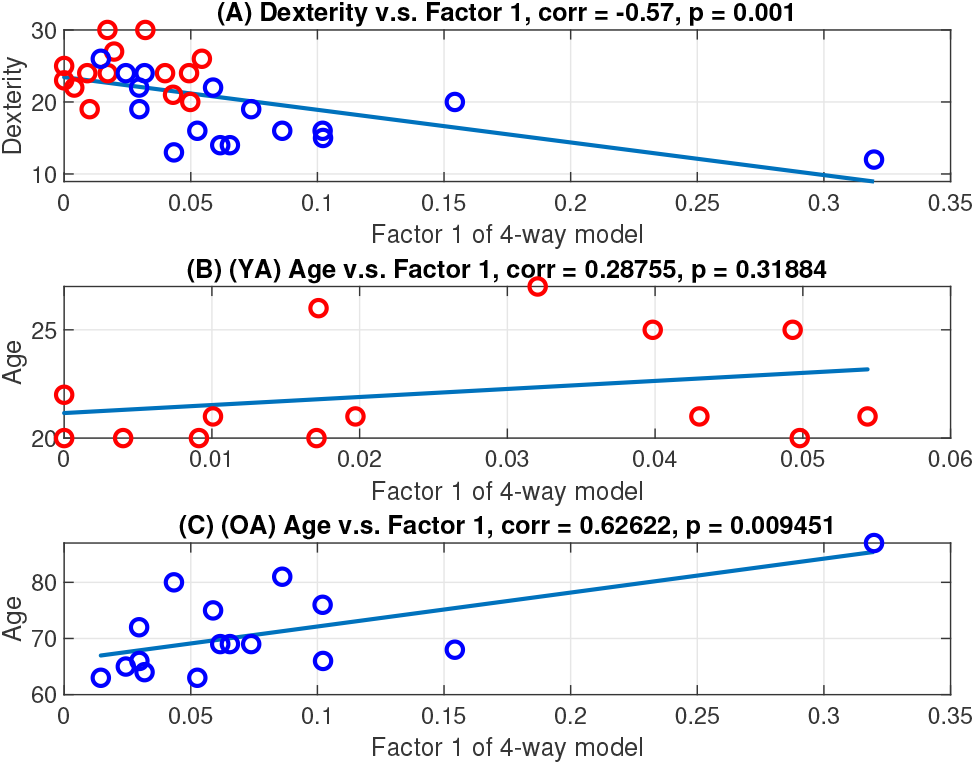
Correlation analysis results between factor 1 and PP or age. (A) Scatter plot showing the participant factor 1 versus PP (manual dexterity index) for all participants. (B) & (C) scatter plot showing the participant factor 1 versus age for the YA & OA group respectively. For all of the above figures, each dot represents a different participant (red for YA and blue for OA, respectively) and the blue line is the linear regression line that fits the data. Figure titles indicate the computed correlation values and the corresponding p-values.

We then computed the correlation between the participant factors and age. Since we had two distinct age groups (YA where age<30 and OA where age>60), we computed correlations for the YA/OA groups separately. For the YA group, the first component had a positive but not significant correlation with age (*rho* = 0.288, *p* = 0.32, Figure 7B). However, for the OA group, there was a significantly positive (*rho* = 0.63, *p* = 0.009) correlation between age and the first factor indicating a higher activation of the first component for older people (Figure 7C). Overall, our findings show higher recruitment of the first component for participants with poor manual dexterity and advanced age.

## Discussion

In this study, we introduced a tensor decomposition and applied it to muscle activation and kinetic signals to extract the motor patterns that underpin object grasping and lifting and investigate any changes induced by ageing. This approach provided a succinct characterisation of the muscle couplings and the related grip and load forces together with the corresponding temporal profiles during the grasp-to-lift movements. Crucially, we extracted these patterns across participants, thus we could characterise the differences in motor patterns attributed to ageing and/or motor decline.

Specifically, we identified two components that activate the same set of muscles in different proportions thus producing different GF to LF ratios. Both components involve the ECR and APB muscles which are crucial for grasping and lifting objects. The first component activates ECR and APB synergistically and shows an increasing-first decreasing-later temporal profile, overall associated with higher GFs compared to LFs. This component is more prevalent in OAs compared to YAs and shows a positive correlation with age and a negative correlation with manual dexterity. The second component instead has a relatively higher activation of APB and a decreasing temporal profile and is associated with a more economic application of GF relative to LF. This component is more prevalent in YAs compared to OAs. Taken together, our findings suggest that ageing results in a reduced capacity to individually activate the muscles in the forearm (ECR) and hand (APB) which may explain the less economic grasping strategy observed in OAs [5]. This effect correlates strongly with a descrease in manual dexterity thus suggesting that the coupling between ECR and APB and higher GF/LF may be a compensatory mechanism for poor manual dexterity as a result of ageing [16].

The presence of the same muscles in both components also has implications about the effect of ageing on muscle synergies [34]. Our findings suggest that, although muscle recruitment remains intact with ageing, the balance of these muscle activations is altered thus leading to differential grip force generation. This may imply that ageing may not alter muscle recruitment but the relative composition of muscle synergies. In fact, this relative change may represent a new motor strategy adopted by older individuals to overcome the degradation in manual dexterity [16].

Concerning future use of the proposed approach, we suggest that the methodology developed here can be applied to a variety of motor behavioral experiments. Crucially, our approach enabled the joint analysis EMG and kinetic measurements and consequently the identification of relationships between GF/LF ratios and muscle couplings which would not be directly observable with separate analyses of the two types of measurements (see also [48, 2, 12] for similar approaches in lower dimensions). Thus, this methodology will be useful when attempting to merge information and identify dependencies between different types of motor signals, such as neuromuscular, kinematic and kinetic recordings.

When the recorded motor signals contain responses across multiple locations (e.g., muscles or joints), times (e.g., different phases of movement execution), trials (repetitions of the same task), experimental conditions (e.g., varying distance, speed or load) and participants (with different age, gender and anthropometric measurements), they are naturally expressed as a 5-way tensor. The methodology we presented here decomposes such tensors into combinations of factors each of which has a spatial component (describing which muscles/forces are activated together), a temporal component (describing the temporal activation profile of the spatial pattern), a trial component (describing the level of recruitment of each spatiotemporal pattern in a given trial), a condition component (describing how much each pattern is used on each experimental condition – object weight here), and a participant component (describing the strength of each pattern activation for each participant). We also proposed a set of measures to evaluate the effectiveness of such a decomposition in both approximating the original recordings (VAF) and conveying information about differences across individuals (classification, correlation and regression analyses) [14, 10, 24].

We contend that such a decomposition can be particularly effective when the aim is to: a) tease apart motor patterns with different functional roles [13]; b) reveal their spatial and temporal representations[11]; c) quantify their relative contribution to discrimination between experimental conditions [15, 14]; and d) assess how these patterns may differ across individuals or populations with common characteristics. The latter was the main aim of this study which revealed the muscle couplings and corresponding forces that predict age and/or manual dexterity differences across participants. In future work, differences in tensor structure can be used to test specific hypotheses about how motor signals differ between healthy and impaired individuals (e.g., stroke or spinal cord injury patients) and potentially inform rehabilitation or treatment strategies [43, 39, 25].

## Material and Methods

### Participants

Human participants of two different age groups took part in the study, 15 YA (>60 years old, three left-handed, M = 22.2 ± 2.59 yrs old; F = 14) and 15 OA (<30 years old, two left-handed, M =70.8 ± 7.42 yrs old; F = 11). All participants (*P* = 30) had no known musculoskeletal or neurological conditions and normal or corrected vision. This research was approved by the Research Ethics Committee of the Faculty of Biological Sciences of University of Leeds.

### Clinical tests

Before the experimental session, we performed two clinical tests to assess the participants’ tactile sensitivity and manual dexterity. We used the Semmes-Weinstein monofilament test [52] as a measure of cutaneous sensitivity. We tested eight sites (four per hand) on the participants’ hands (Semmes-Weinstein location reference): middle fingertip (31), index fingertip (21U & 21R) and thumb tip (11). We also quantified manual dexterity using the Purdue Pegboard test. Participants performed the test bimanually to obtain an overall measure of their manual dexterity.

### Apparatus

To perform the grasping tasks, we built two manipulanda made from carbon-filled nylon (width: 40 mm, height: 110 mm, depth: 50 mm) and containing 50 N load cells (Omega, LCM201-50) which enabled recording grip forces. Grip force data were acquired using a 16-bit data acquisition card (National Instruments, USB-6002) and processed using a custom-built program in Labview (v.14). Reliability of recordings was ensured by prior validation tests (< 1% error for forces between 1 and 36N). To track the object kinematics (200Hz sampling frequency), we attached four Qualisys markers to each manipulandum and used Qualisys camera setups (12 cameras in the lab, 5 in the community centres). For both setups, calibration was successful according to guidelines (error < 1.0mm).

### Grasping and lifting task

All participants sat on a chair in front of a table [27]. The table surface was level with their navel and their feet were flat on the ground and the two manipulanda were placed 75% of shoulder width and 70% of maximum reach for each participant. The participants’ fingers and thumbs were cleaned with alcohol wipes. The researcher demonstrated how to pick up the manipulanda using the two circular plungers, with a precision grip. Participant instructions were to “grasp the objects and lift them level with a target height placed in front of them (300mm height), and to hold the objects as still as possible” [27]. After a 10-second period, the researcher asked the participants to replace the object(s) back on the starting markers. Participants performed *K* = 10 consecutive repetitions of bimanual grasp-lift-replace movements with *J* = 2 object masses: light (200g) and heavy (400g).

### EMG data collection

EMG data were collected from the Anterior Deltoid (Ant Del), Flexor Carpi Radialis (FCR), Extensor Carpi Radialis (ECR) and Abductor Pollicis Brevis (APB) of both arms and hands using eight Delsys Trigno(tm) sensors (sampling frequency of 2,000Hz).

### Data pre-processing

Six degree-of-freedom models were created in Qualisys for each object and were used to compute the position (*x, y, z*), velocity and acceleration of each object. Net load force was calculated from the objects’ mass times a product of the net acceleration [36][23][22]. Finally, to obtain smooth force profiles, grip force (GF) and load force (LF) recordings were low-pass filtered (12Hz cut-off, 4th-order Butterworth).

The EMG recordings for each trial were digitally, full-wave rectified, low-pass filtered (10Hz cut-off, 4th order Butterworth, zero-phase distortion; R Signals package, filtfilt function) and down-sampled to 200Hz to align with kinetic and kinematic datapoints. All trials were visually scanned for artifacts and affected trials were excluded from further analysis (<5% of total number of trials). Then, EMG signals of each participant were normalised against the peak EMG amplitude across all trials, resulting in normalised EMG data values between 0 and 1 [26].

For each trial, EMG, GF and LF data were selected during the dynamic phase of object lifting – from the point of first contact to the end of the object lift / beginning of the stable phase of the object hold. Determination of this time window was done based on the recorded kinematic data of the object position. Specifically, to account for electromechanical delays between EMGs/forces and resulting kinematics, we selected EMG, GF and LF measurements starting 100ms before the first contact with the object and ending 100ms after beginning of the stable phase of object hold.

EMG signals of all muscles, GFs and LFs were time-normalised to *T*_0_ = 500 datapoints to ensure equal temporal weighting across participants and conditions for the subsequent analysis. Then, EMG data and bimanual GFs and LFs for each trial (*K* = 10 for each object *J*) of each participant were concatenated to form a 12 *×* 500 matrix containing all recordings from 8 muscles and 4 forces (*M* = 12) over *T*_0_ = 500 datapoints. Finally, to remove edge artifacts from signal filtering, the first and last 20 samples were removed from each recording, which resulted in signals of *T* = 460 datapoints.

### Tensor formulation and decomposition

The aforementioned data was collated across objects (*J*), trials (*K*), and participants (*P*, ordered according to age from youngest to oldest) to construct the 5-way tensor 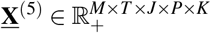, where *M* = 12, *T* = 460, *J* = 2, *P* = 30, and *K* = 10. Averaging the signals across all *K* = 10 trials, a 4-way data tensor 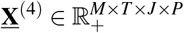 was also constructed for subsequent analyses.

#### Non-negative canonical polyadic (NCP) decomposition

We applied the NCP decomposition to the 5-way EMG data data tensor 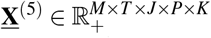. NCP entails a low-rank approximation **<u>X</u>** ∼ **A** ◦ **B** ◦ **C** ◦ **D** ◦ **E** subject to non-negativity constraints on the entries of the so-termed factor matrices {**A, B, C, D, E**}. Non-negativity is well motivated to better interpret the EMG and net load force data. The outer product (◦) decomposition is defined such that scalar tensor entries *X*_*mt jpk*_ are approximated as

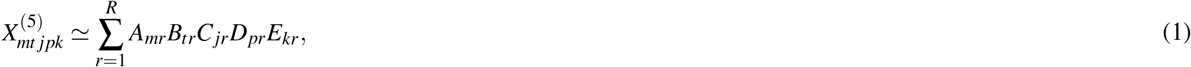

where *R* is the tensor rank adopted for the approximation. Matrix 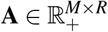 denotes the spatial (force and muscle) factor and its columns 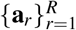 represent the spatial synergies. The row indexing of forces and muscles follows the description in Table 1. Likewise, matrix 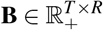 is the temporal factor and its columns 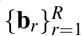 are the temporal activation sequences of each component. Matrices 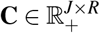 and 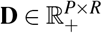 are the object and participant factors, respectively, that capture the object-wise and participant-wise information in the data. Finally, 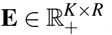 is the trial factor.

Given **<u>X</u>**^(5)^ and a prescribed value of *R* (the method used to choose *R* is described below), the factor matrices are estimated by solving the following non-convex NCP decomposition problem

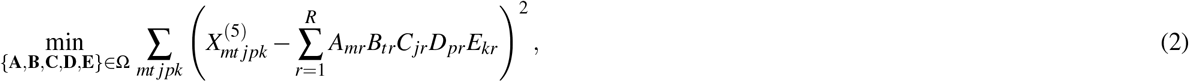

where Ω is the feasible set imposing all optimization variables (i.e., the entries of the factor matrices {**A, B, C, D, E**}) are non-negative. Likewise, when trials are averaged the NCP decomposition is given by **<u>X</u>**^(4)^ ∼ **A** ◦ **B** ◦ **C** ◦ **D** and the factor matrices are obtained by solving

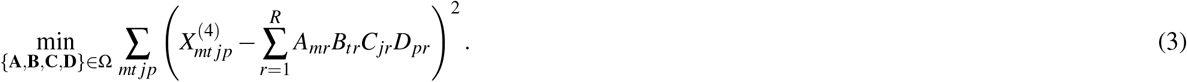

Notice how in the 4-way model there is no trial factor **E**.

Both optimization problems (2) and (3) were solved via the *structured tensor fusion* (SDF) algorithm [46], specifically by using the `sdf_nls` function from Matlab’s toolbox Tensorlab [50]. Multi-way decompositions such as NCPD suffer from an inherent scaling ambiguity. In order to fix the scale, the estimated factors in the 5-way model were normalized by letting ∥**A**∥_*F*_ = ∥**B**∥_*F*_ = ∥**D**∥_*F*_ = ∥**E**∥_*F*_ = 1 and **C** ← ∥**A**∥_*F*_ ∥**B**∥_*F*_ ∥**D**∥_*F*_ ∥**E**∥_*F*_ **C**, where ∥ ·∥_*F*_ denotes the Frobenius norm of its matrix argument. Similarly, ∥**A**∥_*F*_ = ∥**B**∥_*F*_ = ∥**D**∥_*F*_ = 1 and **C** ← ∥**A**∥_*F*_ ∥**B**∥_*F*_ ∥**D**∥_*F*_ **C** for the 4-way model. For *R* = 4 components, the estimated factors for the 5-way and 4-way models are depicted in Figures 2 and 3, respectively.

#### Variance accounted for (VAF) and rank selection

The sole parameter for the NCP decomposition is the rank *R* of the tensor approximant, which naturally affects model complexity as well as the reconstruction error. In order to assess how well the multi-way signal is reconstructed from the estimated synergies, the variance explained by the NCP decomposition is evaluated for different values of *R*. Following a well established criterion, see e.g., [21], the smallest value of *R* is chosen that explains at least 80% of the variance in the original data tensor. To this end, the Variance Accounted For (VAF) metric defined as

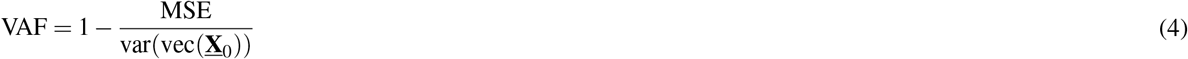

was evaluated, where vec(*·*) stands for tensor vectorization and **<u>X</u>**_0_ denotes the 5-way data tensor **<u>X</u>**^(5)^, or, the 4-way data tensor **<u>X</u>**^(4)^. The mean squared error (MSE) was computed as 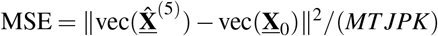 for the 5-way model and 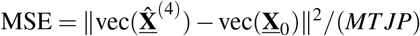 for 4-way model, where 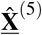 and 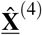 represent the reconstructed tensors for each of the models considered. In other words, the MSE corresponds to the cost functions in the NCP optimization problems (2) and, up to scaling.

Over a grid of candidate values *R* = 1, …, 20, the VAF was computed for both of the 5-way and 4-way tensor model. Results are depicted in Figure 4. With *R* = 4 factors, the obtained NCP decomposition of the 4-way data tensor satisfies the target criterion. The data in the 5-way tensor is noisier because trials are not averaged, hence, as expected, the VAF is smaller than its 4-way counterpart for all values of *R*.

### Correlation analyses

Correlation analyses were conducted to examine if there is any significant relationship between the recruitment of the estimated muscle/force couplings and clinical measures such as manual dexterity and tactile sensitivity, or participant characteristics (age). Specifically, the correlation was computed between the participant mode factors of the identified decomposition (i.e., the columns 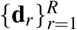 of matrix 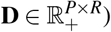) and the *P*×1 vectors of manual dexterity (*PP*) and age information collected across all participants. Bonferroni correction for multiple comparisons was implemented.

### Classifying age groups via linear support vector machine (SVM)

An age classification analysis was conducted to identify any differences in muscle recruitment and GF/LF relationship between the two age groups. Specifically, the columns 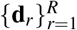 of the participant factor matrix 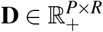 (representing the level of recruitment of each factor for each participant) were used as predictors of the participant age group (YA vs. OA). To this end, a linear support vector machine (SVM) classifier was adopted and trained using leave-one-out cross-validation, which is appropriate when the sample size is limited. Matlab’s `Classification Learner` app was used for the implementation. An iterative procedure was utilized in order to identify the participant factors (and combination thereof) that were most discriminative of age. First, each individual participant factor was separately used as predictor of age group. This served to determine the most discriminative factors. The predictive power of pairs of factors was subsequently examined (to assess if their combination has more age classification power), then triplets, and so forth. Classification accuracy (% of correctly classified participants) and area and the ROC curve (AUC) were adopted as measures of classification performance. Performance did not improve when considering combinations of 3 or more factors.

### Predicting age and manual dexterity via linear regression

Linear regression was utilized to further assess how well manual dexterity and age can be predicted by combinations of the identified factors. The participant factors 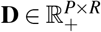 served as predictors and the *P*×1 age vector **y**^(age)^ (or the manual dexterity vector **y**^(*PP*)^) as response variables. The linear regression model 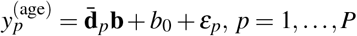, was implemented, where 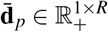 denotes the *p*-th row of **D** and *ε*_*p*_ is a zero-mean noise. Regression coefficients **b** ∈ ℝ^*R*^ (and the corresponding p-values) were estimated using the `fitlm` function in Matlab, and for each predictor these coefficients represent the age- or dexterity-predictive power of each factor. The linear regression results are shown in Figure 6.

## Author Contributions

W.S., R.O.C., S.L.A. conceived the experiments, W.S. conducted the experiments, C.Y., S.S.S., G.M., I.D. analysed the data. G.M., I.D. supervised the analysis and provided conceptual advice. C.Y., S.S.S., G.M., I.D. wrote the manuscript. All authors commented on the manuscript.

